# SARS-CoV-2 minor variant genomes at the start of the pandemic contained markers of VoCs

**DOI:** 10.1101/2022.06.10.495670

**Authors:** Xiaofeng Dong, Julian A. Hiscox

## Abstract

SARS-CoV-2 emerged through limited zoonotic spillovers and was predicted to have constrained sequence diversity. The dominant consensus and minor variant genomes were determined from the earliest samples associated with the Huanan market and the start of the pandemic. The sequence data confirmed that the dominant consensus genomes shared very close homology. However, there were minor variant genomes present in each sample, which encompassed synonymous and non-synonymous changes. Fusion sequences characteristic of defective RNAs were identified that could be linked between patients. Several substitutions (but not deletions) associated with much later variants of concern (VoCs) were already present as minor variant genomes. This suggests it may be possible to predict futures variants at the start of a pandemic by examining where variability in sequence occurs.

---

SARS-CoV-2 is a zoonotic infection that likely had origins in bats and spilled over into the human population via an intermediate animal host. Early cases of the outbreak were associated with the Huanan Wholesale Seafood market (Huanan market) (*1, 2*). Genetic and epidemiological analysis have provided further evidence that the market was the epicenter of the COVID-19 pandemic that resulted in sustained chains of human-to-human transmissions (*3, 4*). The spillover events at the market were likely over a constrained time period with a minimum of two successful spillovers leading to the establishment of transmission of Lineages B and A in the human population, and potentially other spillovers leading to dead ends (*3*). The consequence of this scenario and population bottleneck was limited genetic diversity in the virus at the start of the pandemic in humans. As the pandemic progressed, and SARS-CoV-2 spread through human-to-human transmission both within China and then globally, viral sequences became more and more diverse, with selection pressure acting on those with a fitness advantage including increased transmissibility and immune evasion.

Sequencing information of viral populations within an individual can provide two useful markers. The first is the dominant genome sequence, this is representative of the most abundant sequence in a sample. For patients associated with the Huanan market the dominant SARS-CoV-2 genome sequences were found to be over 99.9% identical with each other (*1, 5, 6*). This was suggested to be indicative of a recent emergence of SARS-CoV-2 into the human population (*5*). The second are the minor genomic variants, which are viral sequences that have a lower abundance than the same site on the dominant genome sequence. The minor variant genomes may contain synonymous and/or non-synonymous (amino acid) substitutions that confer an advantage under selective pressure or affect the viral load and disease phenotype (*7, 8*). Most sequencing studies report and consider the dominant genomic sequence, but hidden underneath, depending on what sequencing approach has been used, is information about the minor variant genomes and therefore the virus population.

To study the minor variant genomes at the start of the COVID-19 pandemic, sequence data from the first cases was analyzed to define the minor variant genome populations. This encompassed samples from patients with a reported association with the Huanan market or earliest samples as reported by symptom onset date or deposition of sequence. We note that this sequencing was not necessarily designed to identify minor variants. Data was identified from 16 patients (Table S1) where SARS-CoV-2 had been sequenced from bronchoalveolar lavage fluid using a meta-transcriptomic approach – in this case Illumina. Different approaches were used to assign minor variants from the sequence data. To distinguish the low frequency variants from sequence errors, DiversiTools was used such that the based calling algorithms used the Illumina quality scores to calculate a P-value for each variant at each nucleotide site (*7*). Alternatively, an arbitrary read depth of 10 or a 100 nucleotides per site was also considered to identify minor variant genomes. We note that coverage across the genome was variable in the sequencing data, with some places having high coverage and some with very low coverage. We acknowledge that coverage at a nucleotide position may influence base calling and confidence in assigning minor variants. As described, the p-value was used to improve the confidence in base calling, given the caveats of low sample read depth in many cases. Ideally this study would be complemented with high read depth sequencing, but we would note this analysis is based on publicly available data, and results should be interpreted with this in mind. For this study patients were ordered by symptom onset date and given a sample ID from S1 to S16 for ease of labeling. How this labeling relates to accession IDs, data deposition and WHO IDs are described in Table S1.

Minor variant genomes were identified across all genes in SARS-CoV-2 for each patient (Fig. 1, for combined patients showing non-synonymous changes and Fig. S1 for individual patients showing non-synonymous changes). The data indicated that for minor variants in some genes amino acid substitutions at specific sites were tolerated, whereas in other genes these were less frequent, including the envelope (E), membrane (M), ORF6, ORF7b and ORF10. At the level of individual patients there were some patients with very little population diversity in minor variant genomes in SARS-CoV-2 including patients S9, S12 and S14 (Fig. S1). There were higher frequency substitutions in SARS-CoV-2 from some patients. For example, in SARS-CoV-2 from patient S6 there were two substitutions (C25R and V49I) in ORF8 that were approximately 40%.

**Fig. 1.**
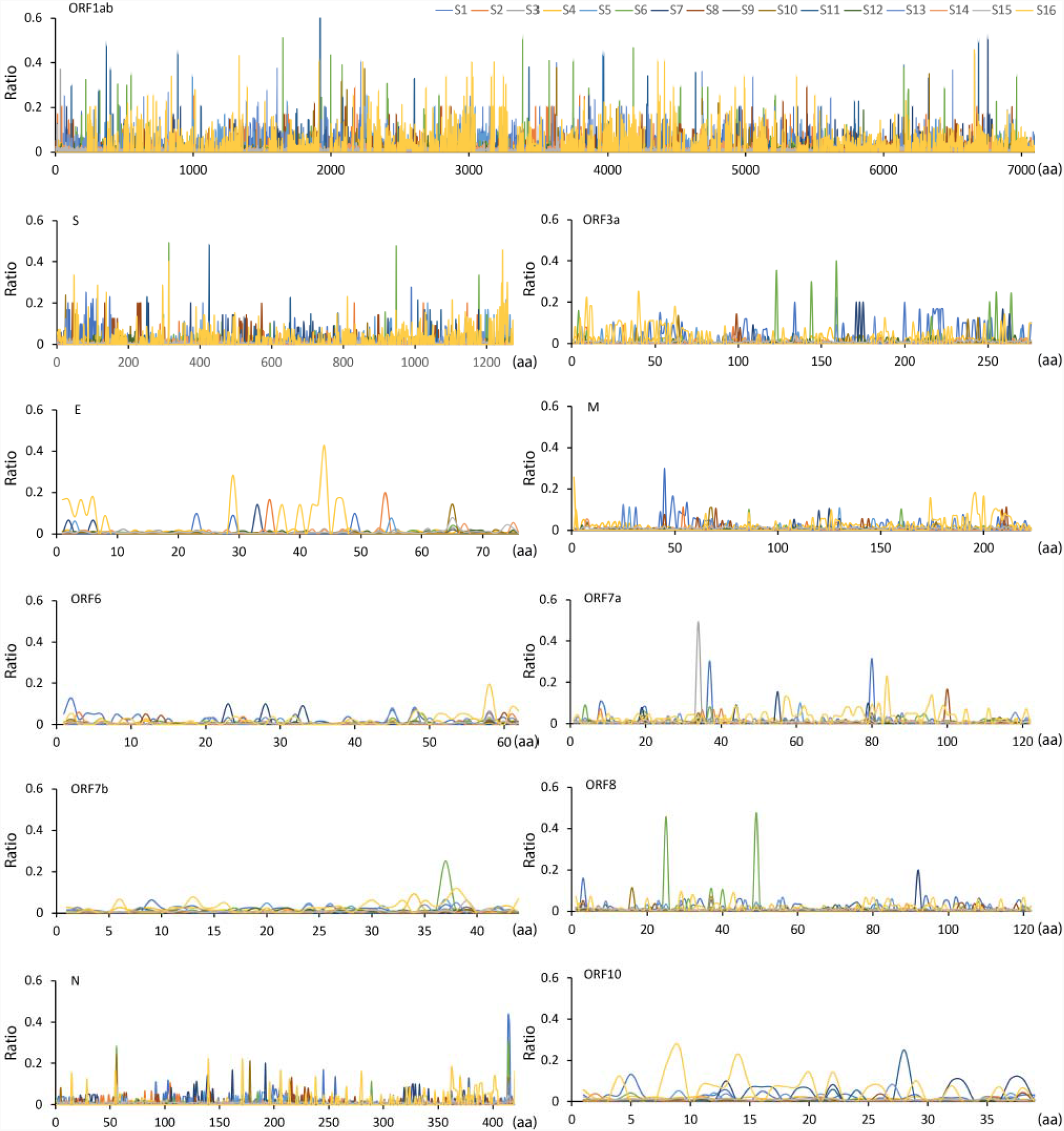
Map of minor variant genomes across the SARS-CoV-2 genome showing non-synonymous substitutions for each gene from samples sequenced from 16 patients down selected for this study. The position of each amino acid is shown on the x-axis and the y-axis shows the ratio of that substitution. Note this cannot be more than 49% otherwise this would be the dominant genome sequence. The amino acid site with coverage >= 5 were shown.

To investigate minor genomic variants and the implications of amino acid substitutions and phenotype, a cut off greater and lower than a 20% threshold, was considered. For a threshold of more than 20% there were four patients that had a greater number of minor variant genomes in SARS-CoV-2, patients S1, S6, S10 and S11 (Fig. 2). Several of the SARS-CoV-2 minor variant genomes sampled from these patients had premature stop codons. For example, premature stop codons were identified in SARS-CoV-2 from patient S11 in NSP2 and in NSP14 and ORF3A from patient S6. As these were present as minor variants the activity of the wildtype protein may have been impacted by a pool of aberrantly functioning protein (*7*).

**Fig. 2.**
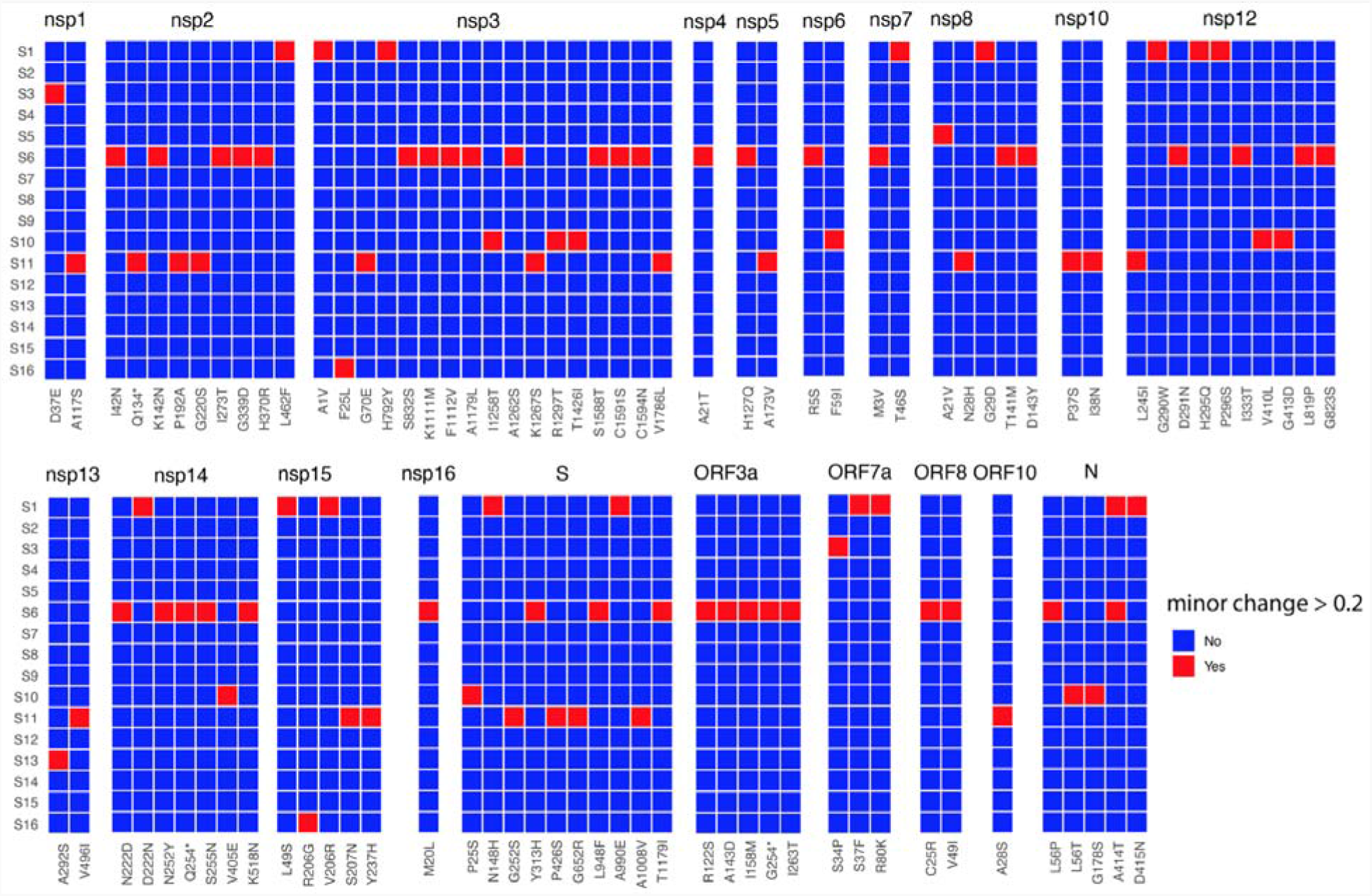
Heat map of non-synonymous changes at the minor variant level in SARS-CoV-2 that have a threshold between 20 and 49% in the 16 patients (y-axis). The panel is divided into each of the SARS-CoV-2 proteins and substitutions are shown on the x-axis. The amino acid site with coverage >= 10 are shown.

Interestingly, several minor genomic variants were identified in the spike protein (Fig. 3A, using p-value) and other viral proteins (Fig. 3B, using p-value) that were subsequently found in different VoCs. These were not uniform in position or frequency in SARS-CoV-2 sampled between the different patients. Nevertheless, the data indicated that in terms of amino acid substitution, the hallmarks of the WHO VoCs (https://covariants.org/variants), were already present at the start of the pandemic or indicated tolerability of substitutions at these positions, in SARS-CoV-2 from some patients. This included the P323L substitution in NSP12 in SARS-CoV-2 minor genomic variants from patients S3, S9, S10 and S15 (Fig 3B). Although below 5% at a minor variant genome level in these patients, data suggests that this substitution is under strong selection pressure and can dominant an infection within days (*9*). Minor genomic variants of SARS-CoV-2 from patient 16 had several substitutions between a frequency of 5 and 15%. These included spike protein substitutions; K417N associated with the ‘Delta plus’ variant; T478K associated with Delta; Q498R associated with Omicron; D614G first associated with an increase in transmission from the Wuhan reference sequence/virus (*10*) and N679K associated with Gamma and Omicron which adds to the polybasic nature of the furin cleavage site. This analysis was based on using p-value to exclude sequencing errors with a pipeline used for identifying minor variant genomes in Ebola virus (*7*) instead of applying a coverage cutoff. However, using a coverage of either at least ten or 100 (Fig. S2 and S3, respectively) some of the substitutions that were assigned based on p-value were not identified in some samples due to the relatively lower sequencing depth.

**Fig. 3.**
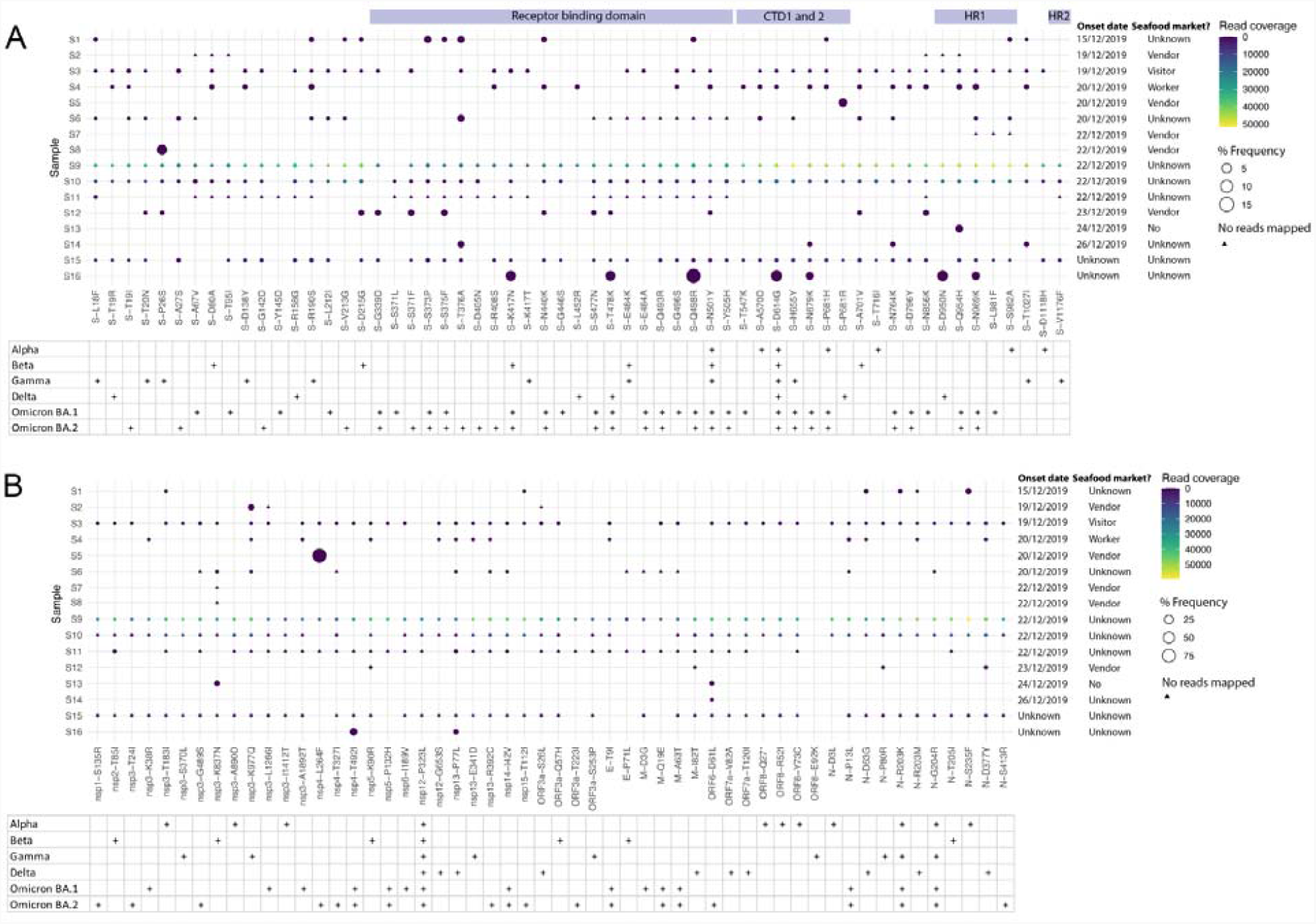
Non-synonymous substitutions in the minor genomic variants of SARS-CoV-2 from each of the 16 down selected patients focusing on sites that define VoCs (https://covariants.org/variants) in the spike protein (A) and other regions of the genome (B). Indicated for each protein is the substitution (x-axis) and its position as well the read coverage and frequency within the viral population for each patient. Features of interest in the spike protein are indicated. Shown are also which VoC the substitution is associated with. Illumina sequence errors were removed as described in methods.

The minor variant genomes may provide the possibility to establish potential transmission chains between the different patients. For example, minor variant genomes of SARS-CoV-2 from patients S3, S9 and S10 appeared to have a similar profile of non-synonymous substitutions suggesting a relationship between them (although we note these were low frequency revealed by high read depth). To investigate these further, common fusion sites were identified in the population of SARS-CoV-2 in each patient (Fig. 4). The formation of fusion sites between disparate parts of the genome is common in coronaviruses due to the high rate of recombination inside a coronavirus infected cell (*11*), and the mechanism of subgenomic mRNA synthesis (*12*). This process results in the formation of defective RNAs. Subgenomic mRNAs were computationally identified and removed from this analysis of the sequencing data (*13*). The deletion analysis pulled out two interesting aspects. The first is that patients S9, S10 and S3 had fusion sites in common. Between S9 and S10 there were three sites in common with fusions between nucleotides 2273 to 2307, 12076 to 12329 and 23795 to 23828. Between patients S3 and S9 there was one site in common between nucleotides 23554 to 23583. This would suggest a potential common source between them. The second was that the fusion sites were most common within the nucleoprotein gene, and between this and other genes. Loss of nucleoprotein is unlikely to be tolerated due to the functions in virus replication (*14*). Amino acid deletions associated with VoCs were not found in these samples, suggesting that these developed much later than the potential amino acid substitutions (Table S2).

**Fig. 4.**
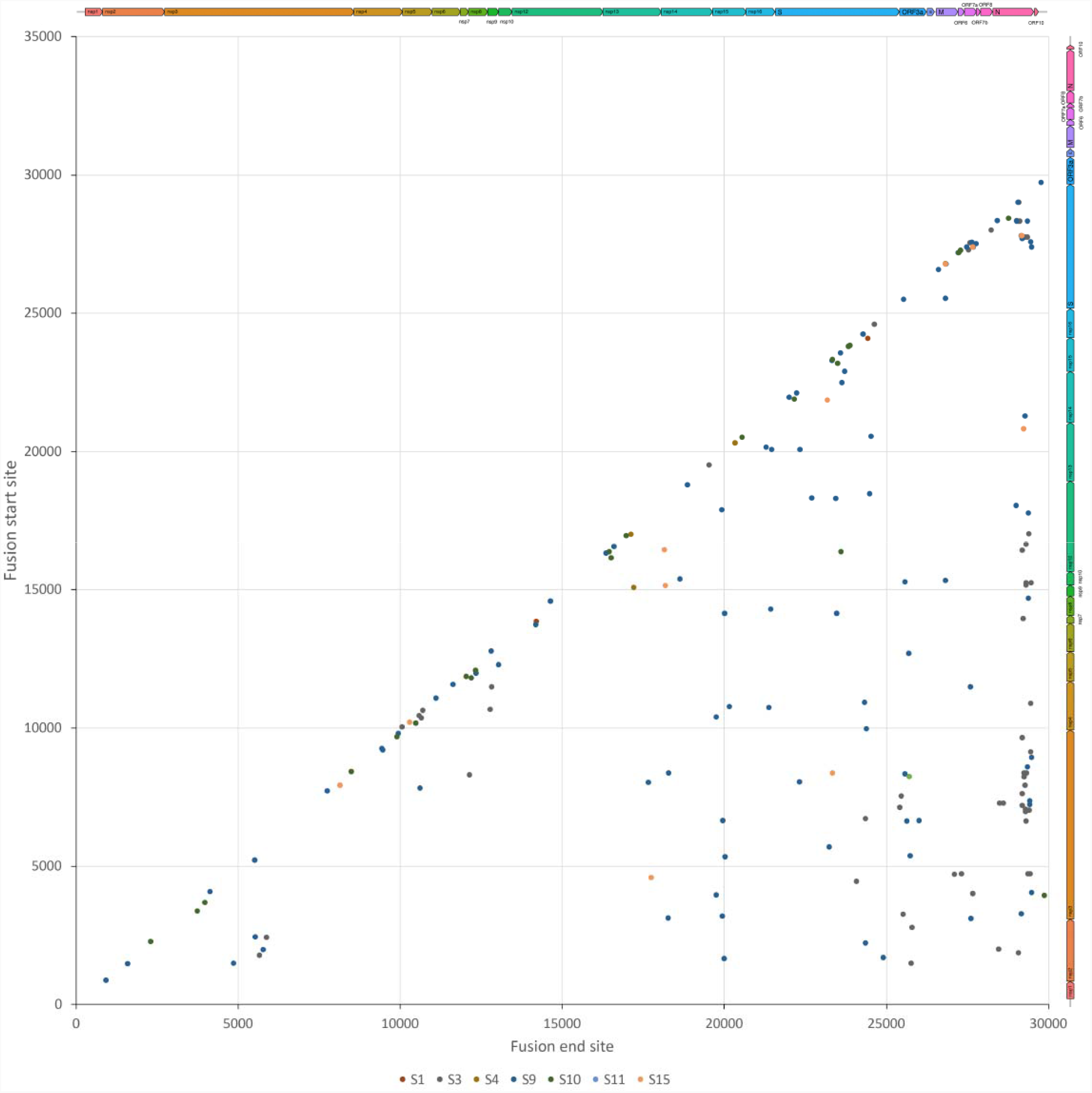
Virus genome position of the start of the fusion site (y-axis) plotted against the end fusion site (x-axis) to show the fusion events along the SARS-CoV-2 genome. Recombination events that generate subgenomic mRNAs as a result of discontinuous transcription during negative strand synthesis have been excluded.

This study has identified potentially interesting sequence features in the SARS-CoV-2 population at the start of the COVID-19 pandemic. Open-source data from samples sequenced by different groups indicated the presence of minor variant genomes. Certain amino acid substitutions within these minor variants were associated with future VoCs. Predicting the emergence of VoCs is a crucial goal in determining their implications for therapeutic treatment regimens and vaccination. Such approaches have in part relied on in vitro evolution studies (*15*) and rapid experimental work once a VoC has emerged. The COVID-19 pandemic and concomitant sequencing of SARS-CoV-2 has provided us with the first extensive guide to the origin and evolution of a virus in real time. This work suggests that analysis of minor variant genomes and the identification of variable sites at the start of viral outbreak can provide a partial playbook for the evolution of the virus. Such information can be used to inform the broad utility of medical countermeasures and challenges to immunity in the face of a virus that has the potential for great diversity.

## Materials and Methods

### Consensus genome and minor variation

The sequencing reads of sample S1-S16 were collected from four published works and NCBI database (Table S1). The consensus genome sequences and minor variations of these samples were generated as our previous description (*16*) (*7*). Hisat2 v2.1.0 (*17*) was used to map the trimmed reads on the human reference genome assembly GRCh38 (release-91) downloaded from the Ensembl FTP site. The unmapped reads were extracted by bam2fastq (v1.1.0) and then mapped on a known SARS-CoV-2 genome (GenBank sequence accession: NC_045512.2) using Bowtie2 v2.4.1 (*17*) by setting the options to parameters “--local -X 2000 --no-mixed”, followed by Sam file to Bam file conversion, sorting, and removal of the reads with a mapping quality score below 11 using SAMtools v1.9 (*18*). After that, the PCR and optical duplicate reads in the bam files were discarded using the MarkDuplicates in the Picard toolkit v2.18.25 (http://broadinstitute.github.io/picard/) with the option of “REMOVE_DUPLICATES=true”. The resultant Bam file was processed by Quasirecomb v1.2 (*19*) to generate a phred-weighted table of nucleotide frequencies which were parsed with a custom perl script to generate a consensus genome sequence (*16*). The consensus genome sequence was then used as a template in the second round of mapping to generate a reference genome sequence for all downstream analysis. Reads (unmapped on human genome) were realigned to the reference SARS-CoV-2 consensus genome sequence using Bowtie2 with the parameter of “--local -X 2000 --no-mixed”. The Bowtie2 outputs were processed in the same way as above to generate a Bam file without read duplication. This Bam file was then processed by diversiutils script in DiversiTools (http://josephhughes.github.io/btctools/) with the “-orfs” function to generate the number of amino acid change caused by the nucleotide deviation at each site in protein. In order to distinguish the low frequency variants from Illumina sequence errors, the diversiutils used the calling algorithms based on the Illumina quality scores to calculate a P-value for each variant at each nucleotide site (*20*). The amino acid change was then filtered based on the P-value (<0.05) by removing the low frequency variants from Illumina sequence errors.

### Insertion, deletion and fusion

Bam file was generated by realigning the reads (unmapped on human genome) to the reference SARS-CoV-2 consensus genome sequence using Bowtie2 with the parameter of “--local -X 2000 --no-mixed”. PCR and optical duplicate reads in the bam file were discarded as described above. Insertion and deletion were then called by analysis of these Bam files using FreeBayes (v1.3.5) (*21*) with the parameter of “--ploidy 1” and quality cutoff of 10.

Sequencing reads (unmapped on human genome) were aligned to the reference SARS-CoV-2 genome (GenBank sequence accession: NC_045512.2) with BWA-MEM (v0.7.17) (*22*). Alignments from BWA-MEM were converted to “bam” format and then analysed by delly (v1.0.3) (*23*) to identify fusion sites on the SARS-CoV-2 genome using “call” function with default setting. Only the fusion events passed all filters of delly were reported.

## Acknowledgments

We would like to thank members of our laboratory for discussions around viral origins and evolution, particularly Rebekah Penrice-Randal, Hannah Goldswain, I’ah Donavan-Banfield and Tessa Prince, and David Matthews at the University of Bristol.

## Funding

United States Food and Drug Administration Medical Countermeasures Initiative contract (75F40120C00085). The article reflects the views of the authors and does not represent the views or policies of the FDA.

Medical Research Council (MR/W005611/1) G2P-UK: A national virology consortium to address phenotypic consequences of SARS-CoV-2 genomic variation.

National Institute for Health Research Health Protection Research Unit (HPRU) in Emerging and Zoonotic Infections at University of Liverpool in partnership with Public Health England (PHE) (now UKHSA), in collaboration with Liverpool School of Tropical Medicine and the University of Oxford (award 200907). The views expressed are those of the authors and not necessarily those of the Department of Health and Social Care or NIHR.

## Author contributions

Conceptualization: JAH

Methodology: XD

Investigation: XD, JAH

Visualization: XD

Funding acquisition: JAH

Project administration: JAH

Supervision: JAH

Writing – original draft: XD, JAH

Writing – review & editing: XD, JAH

