## Supplemental Figures and Tables for "SARS-CoV-2 minor variant genomes at the start of the pandemic contained markers of VoCs"

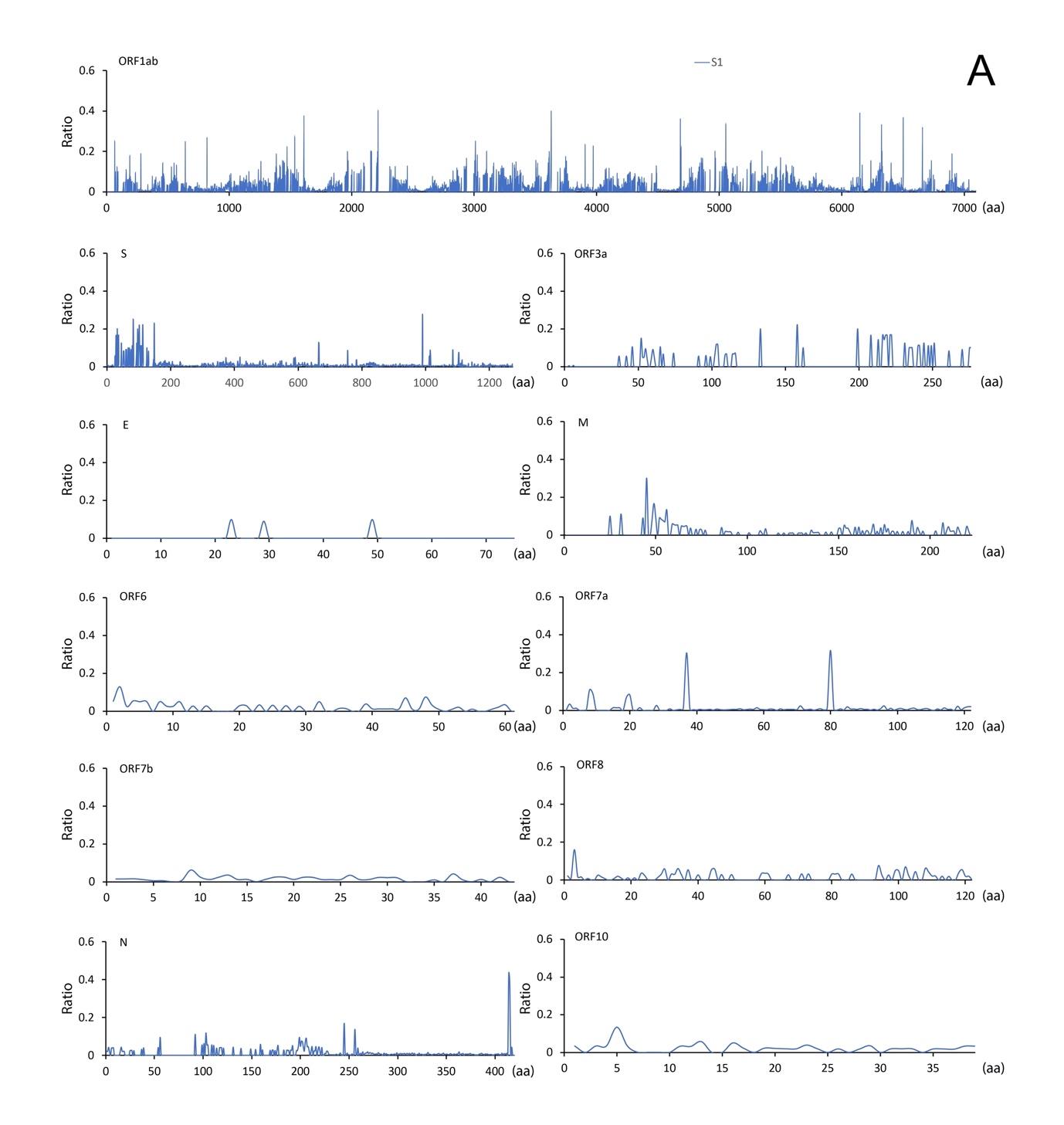

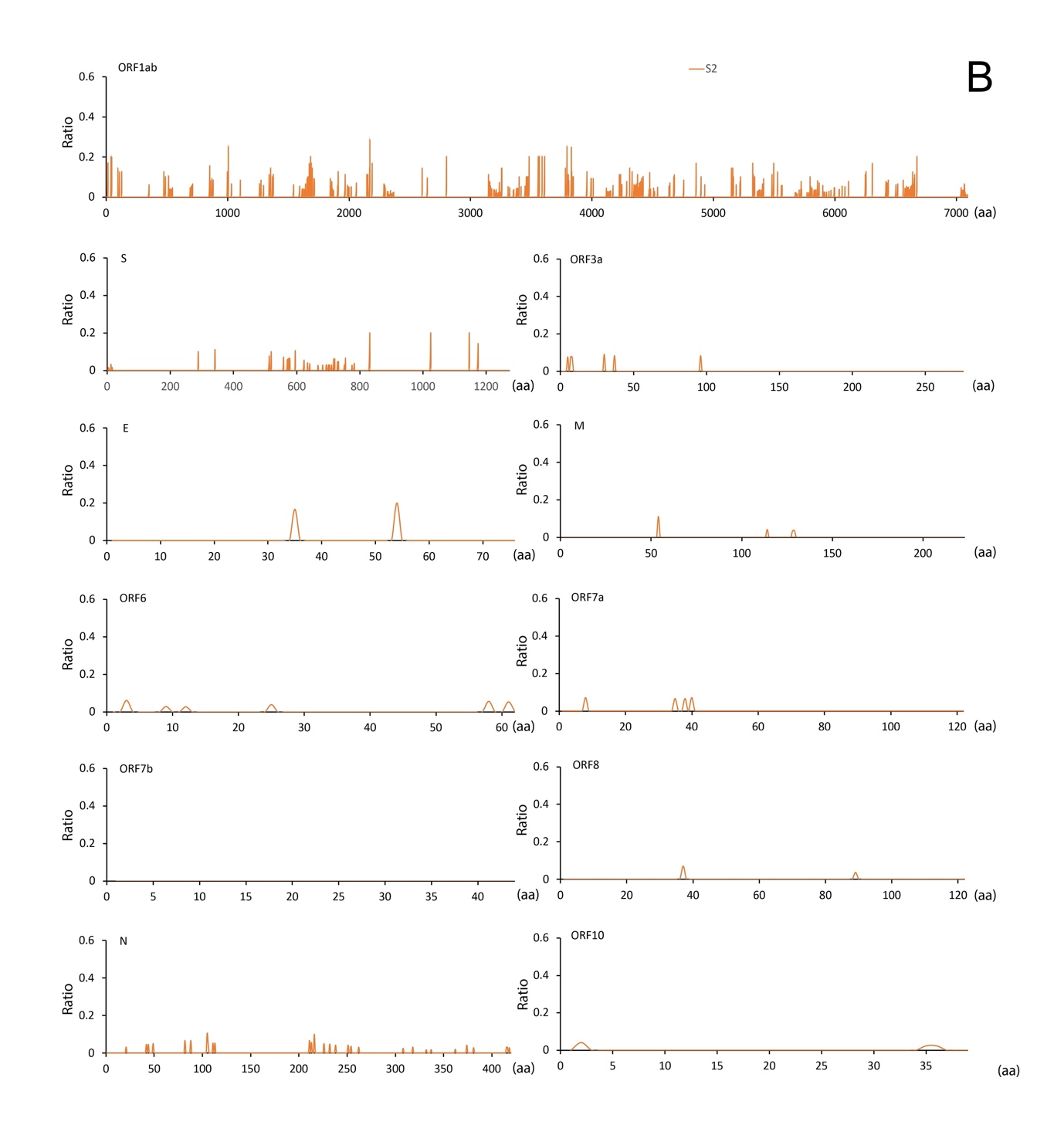

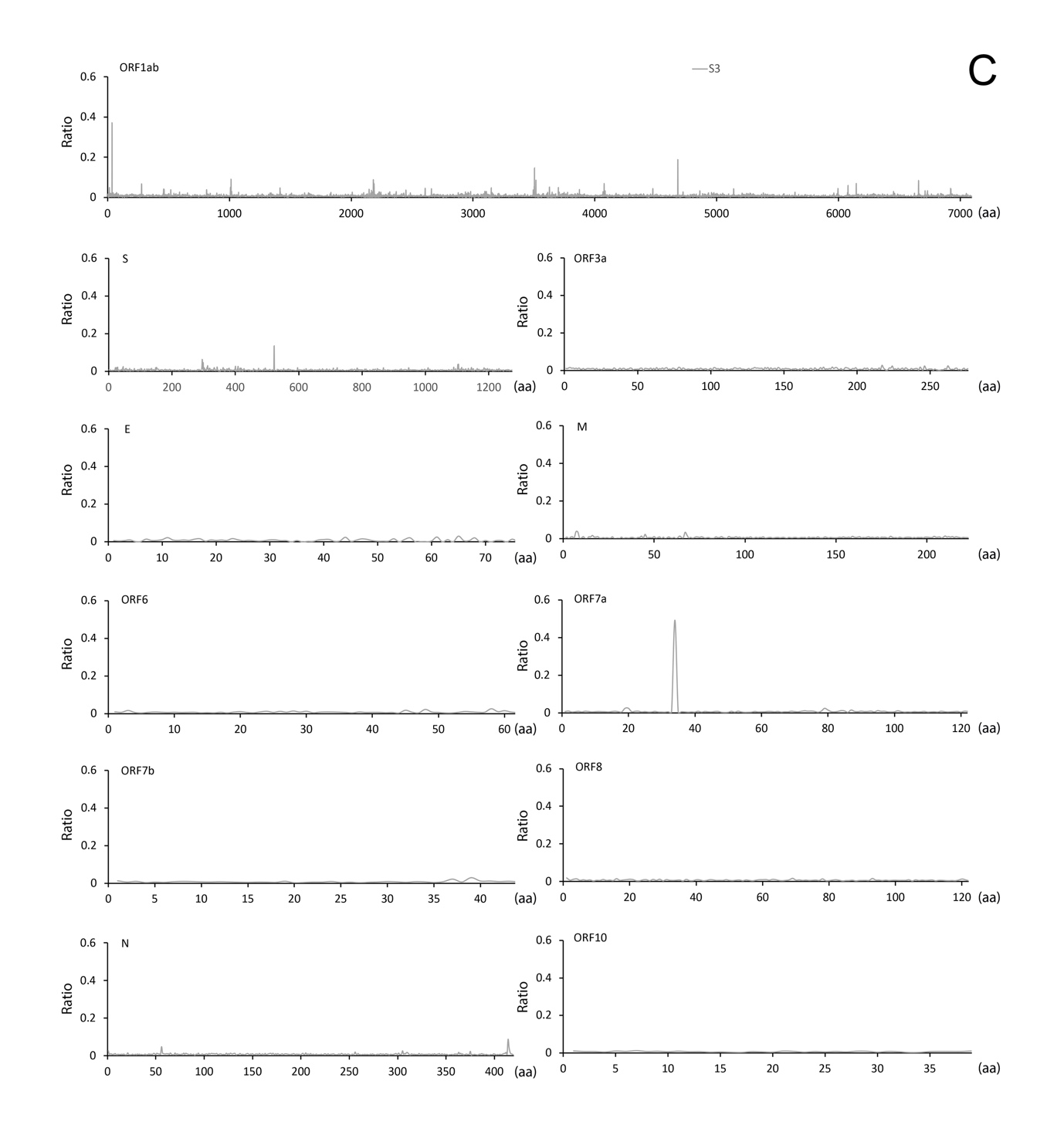

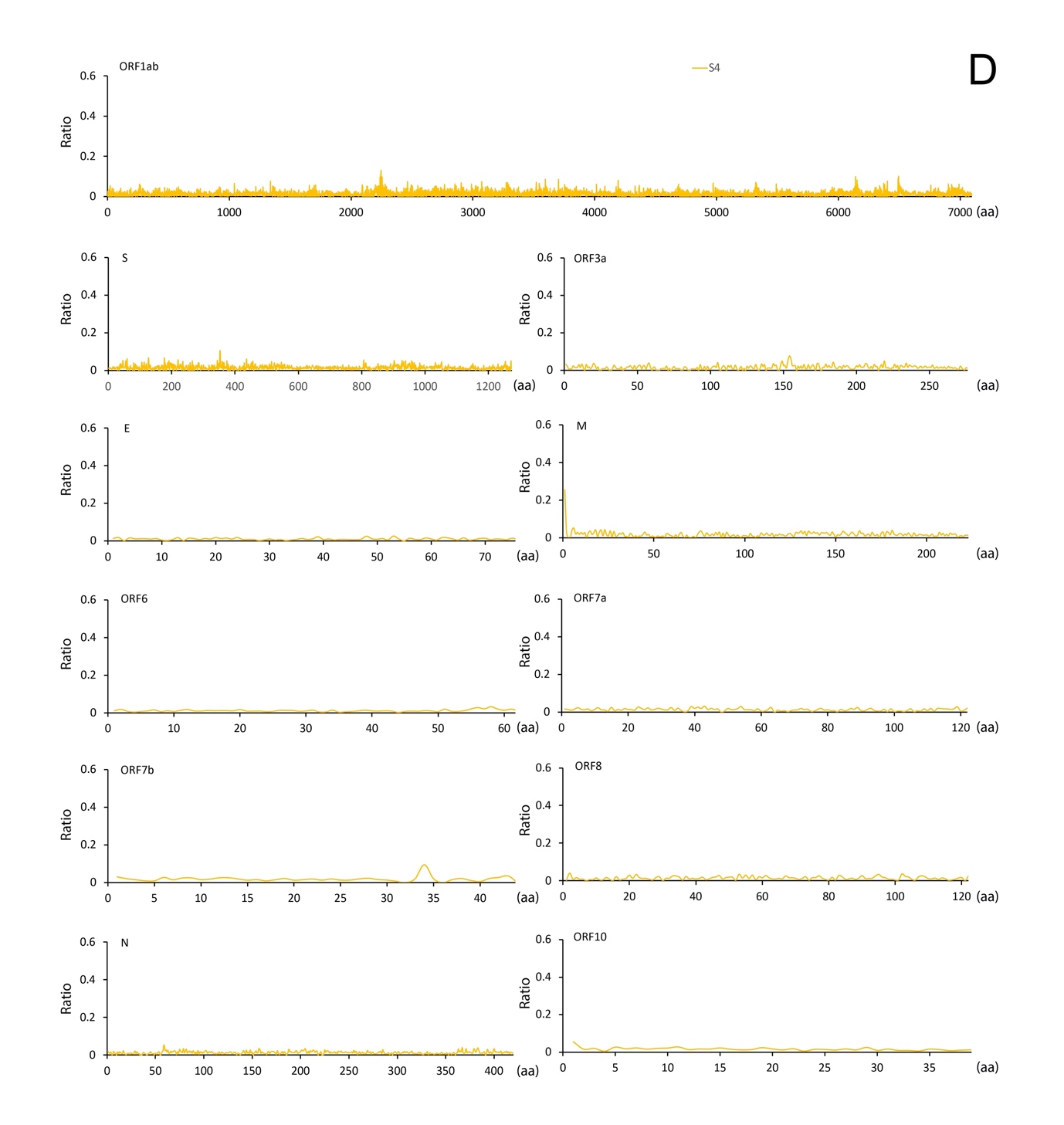

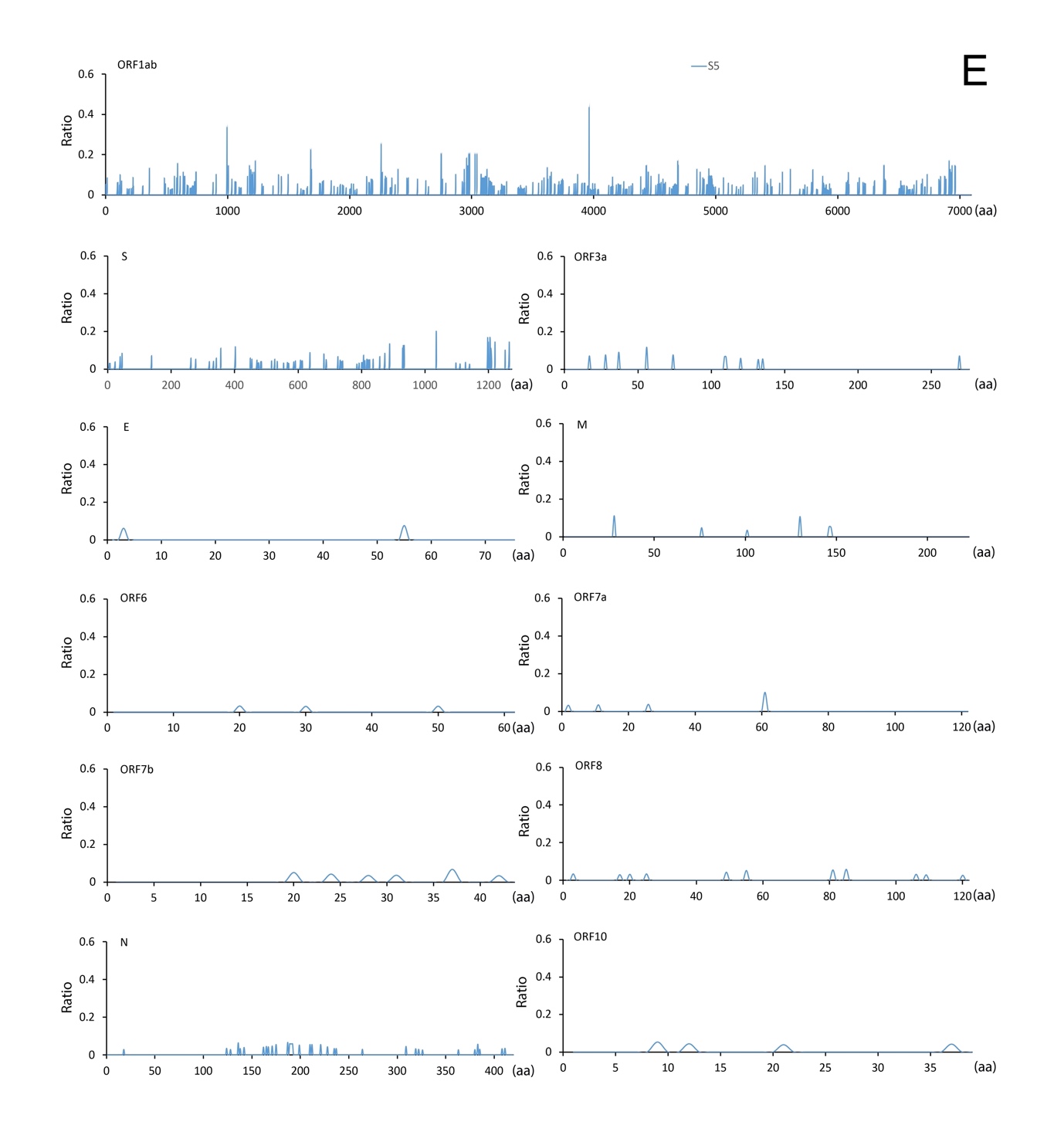

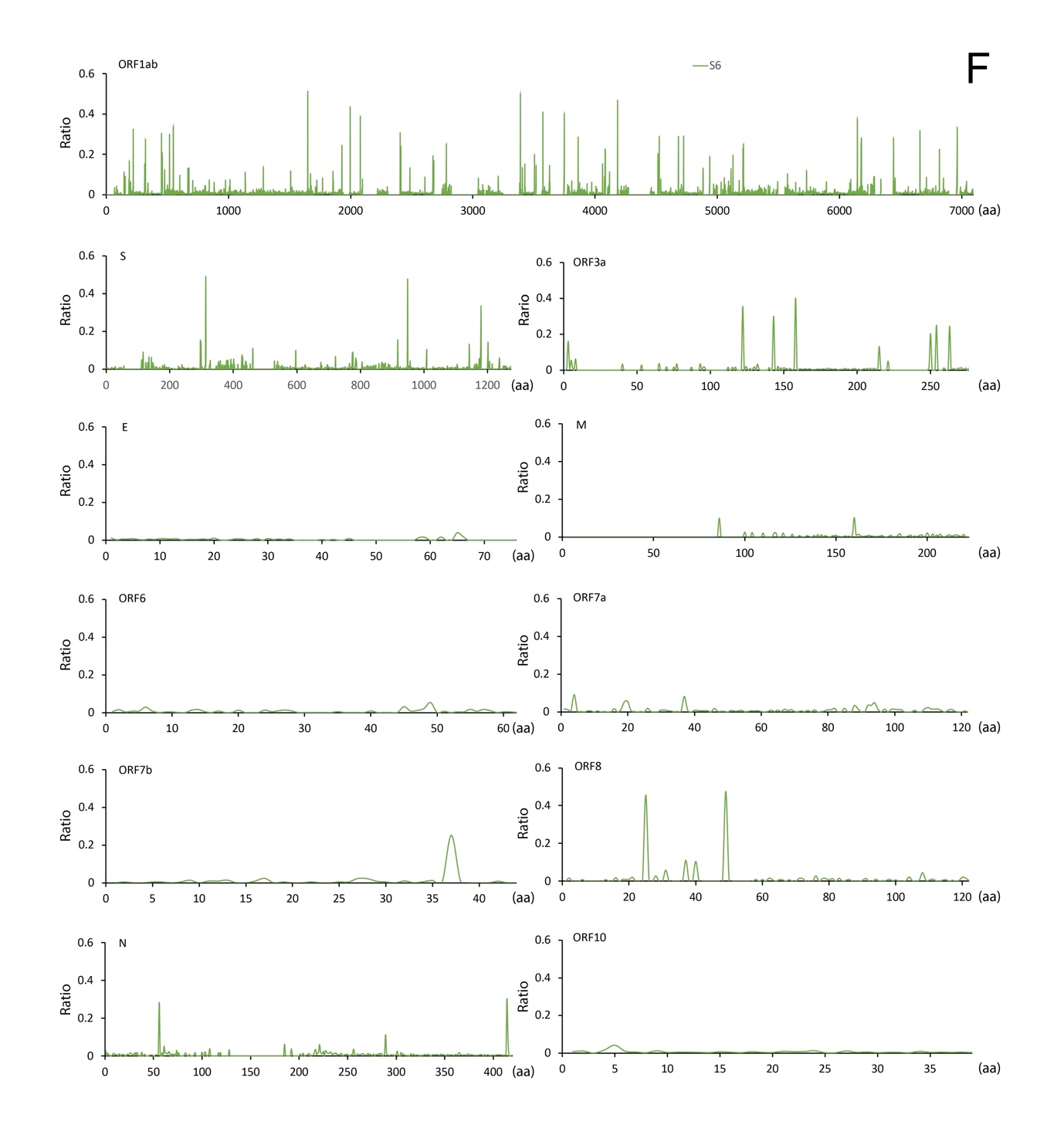

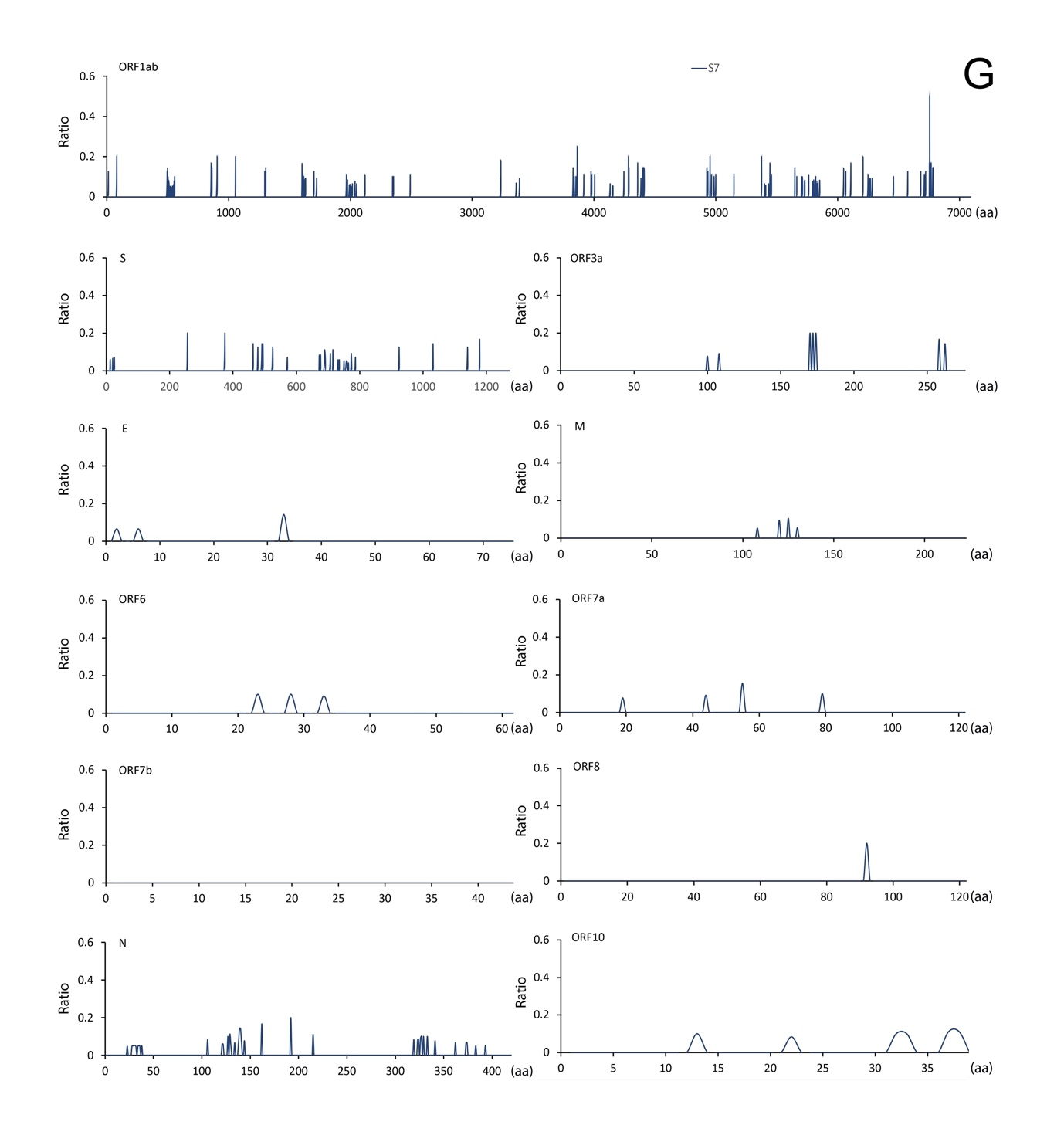

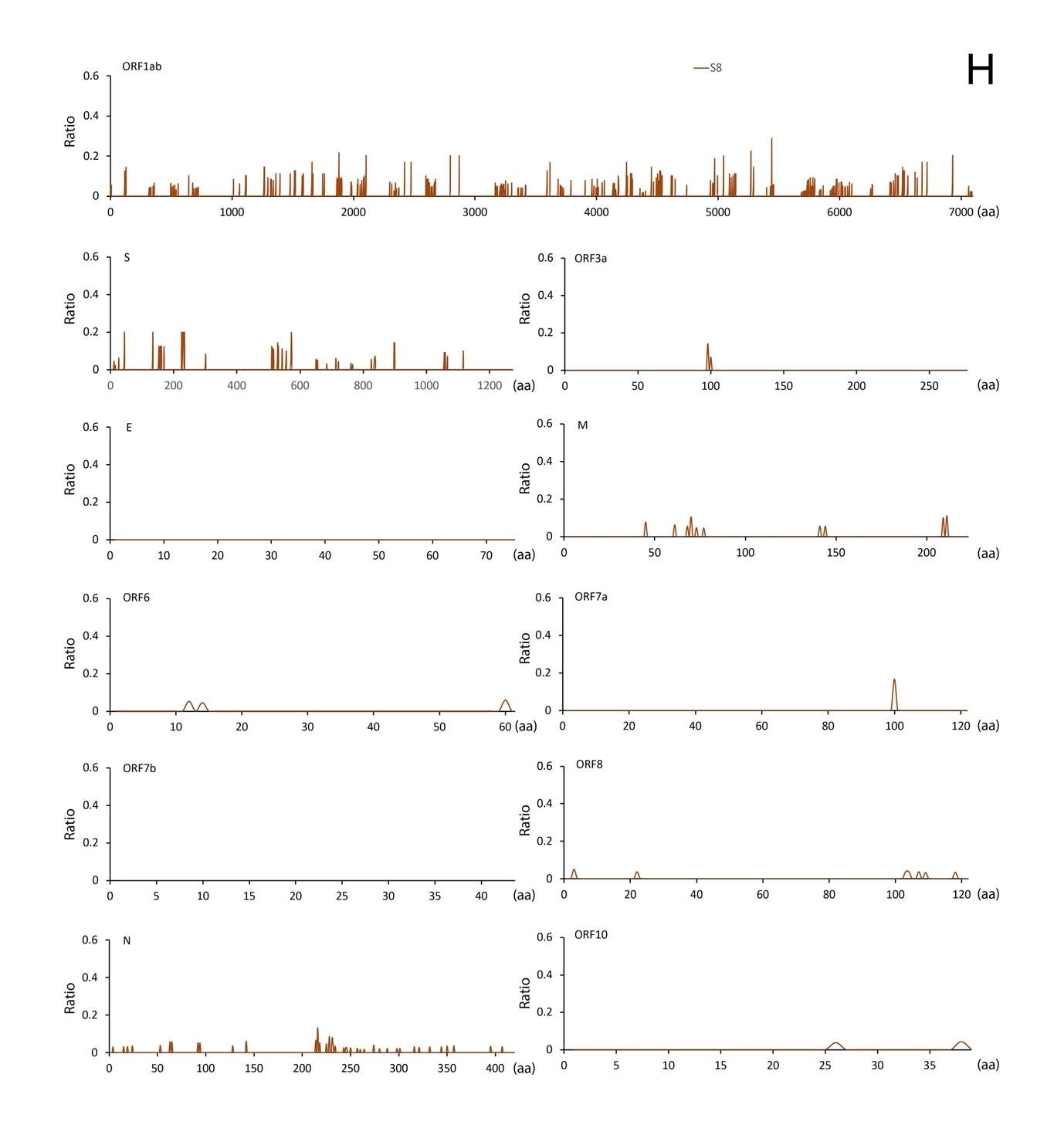

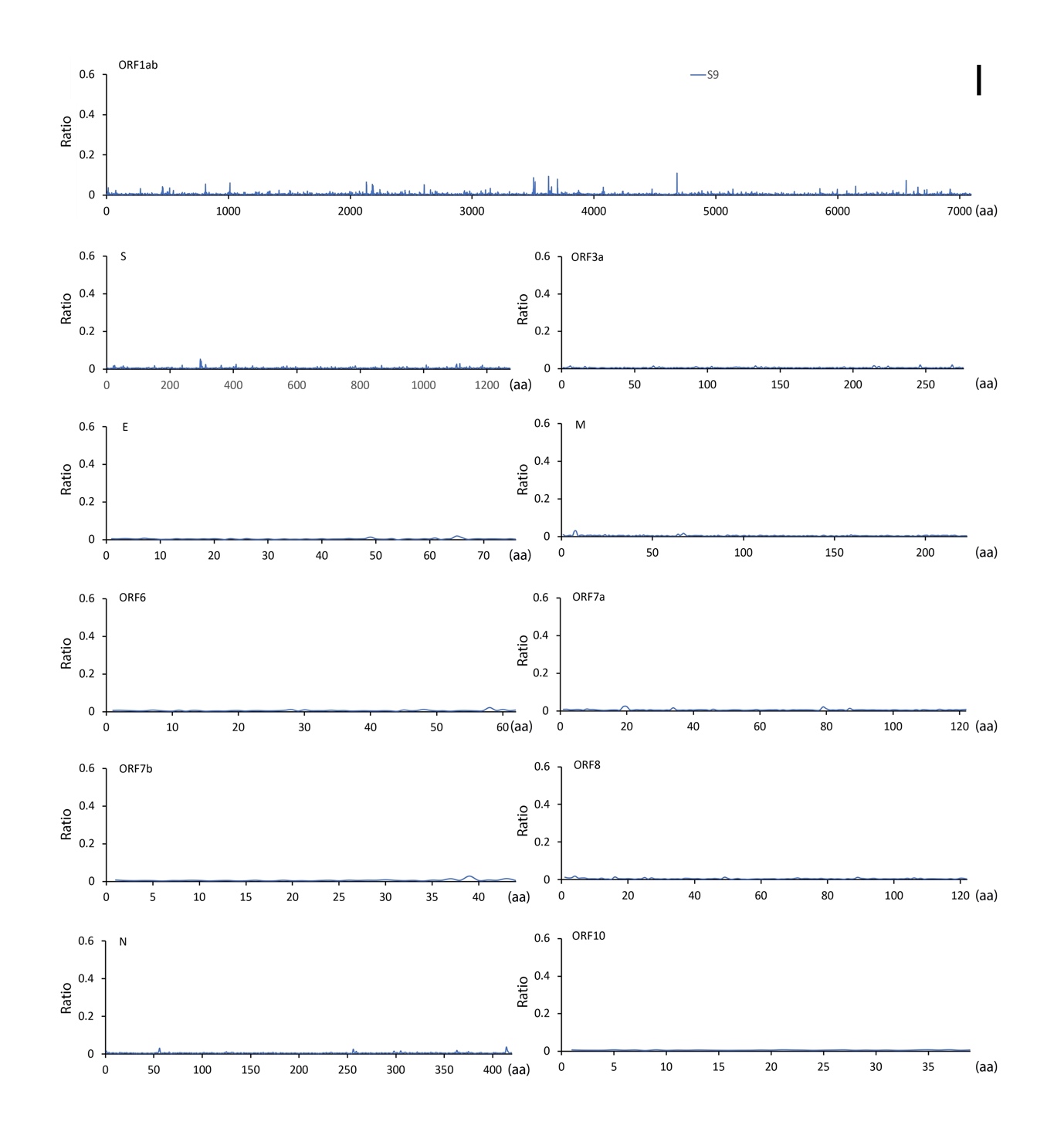

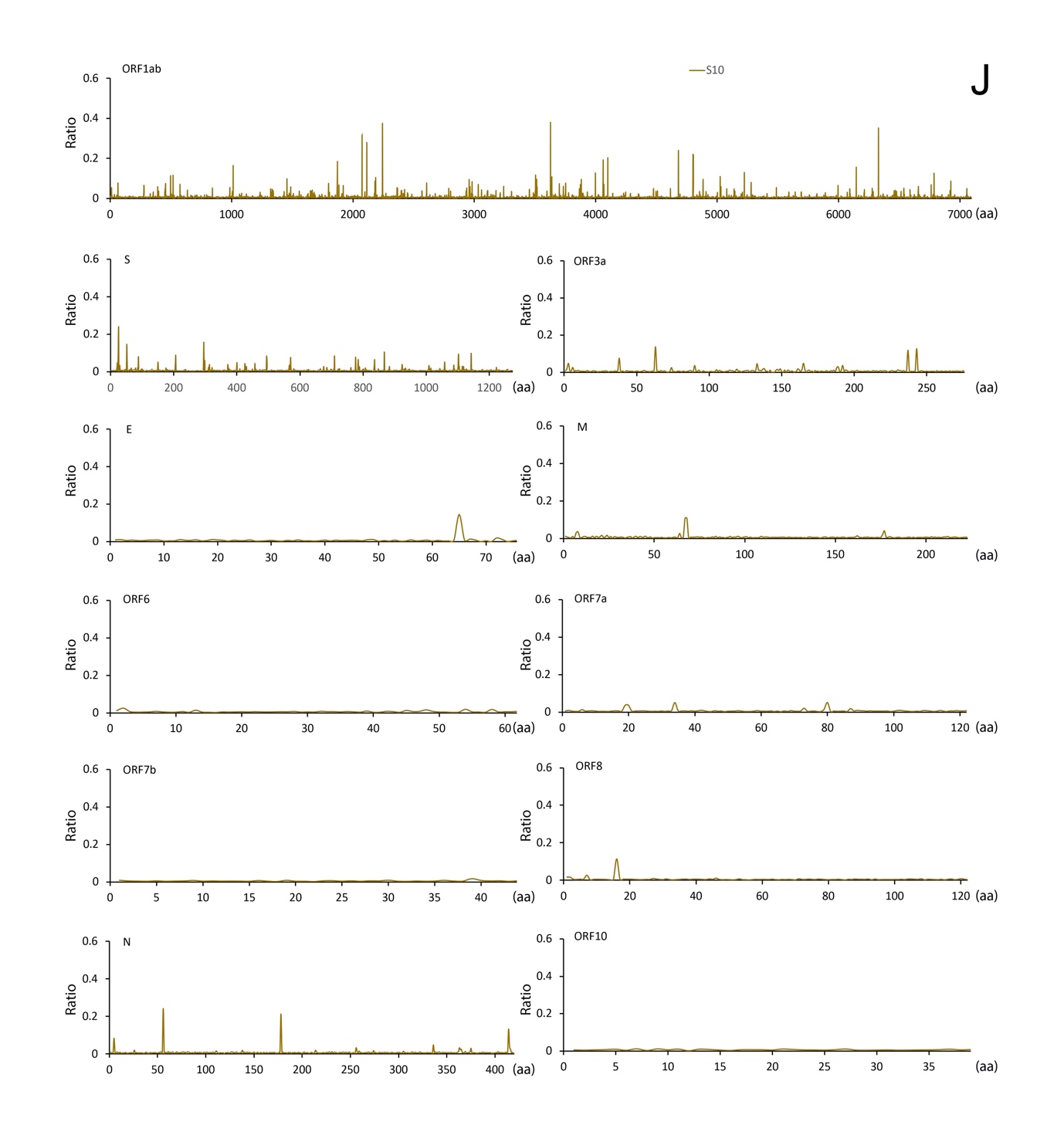

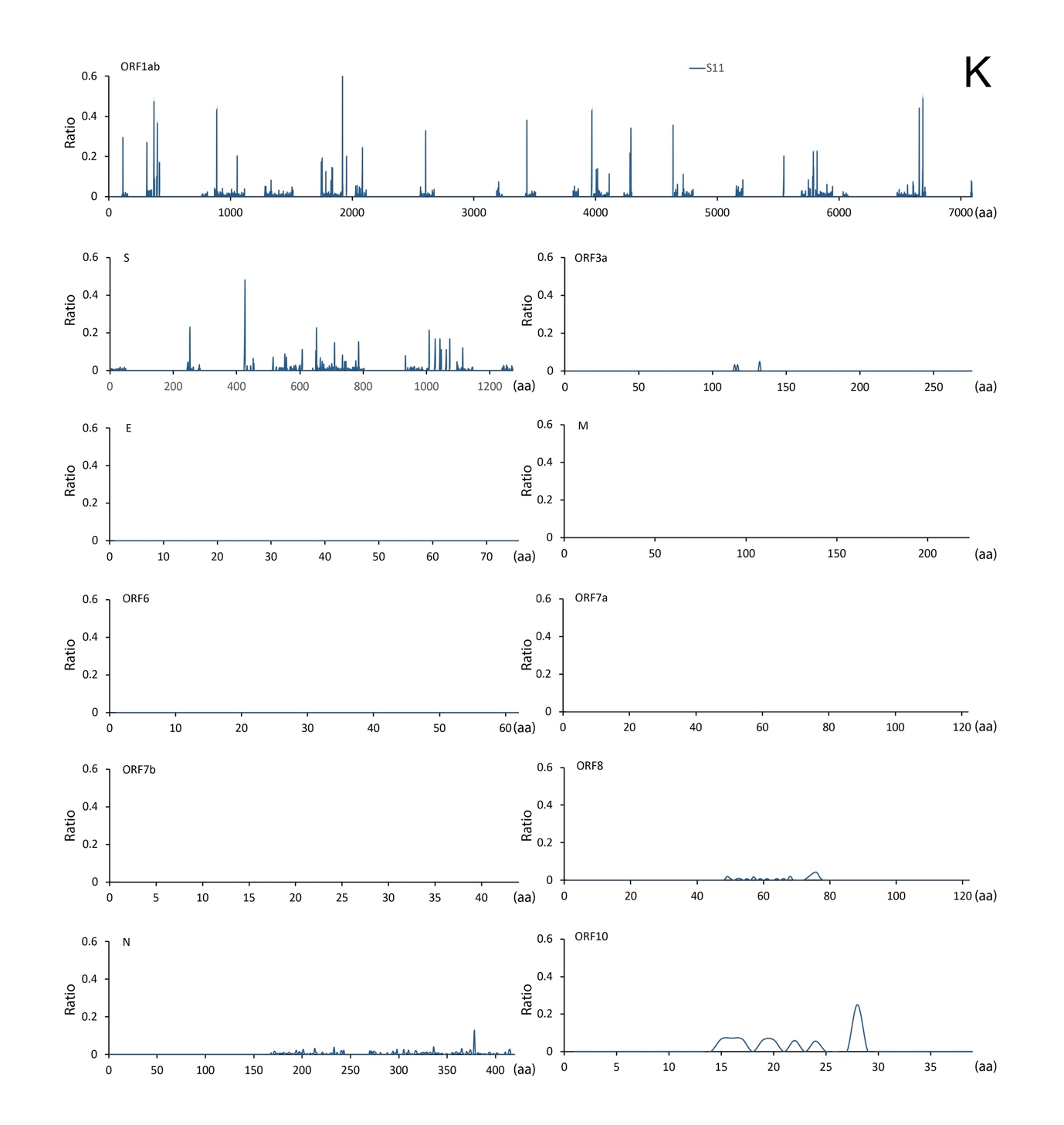

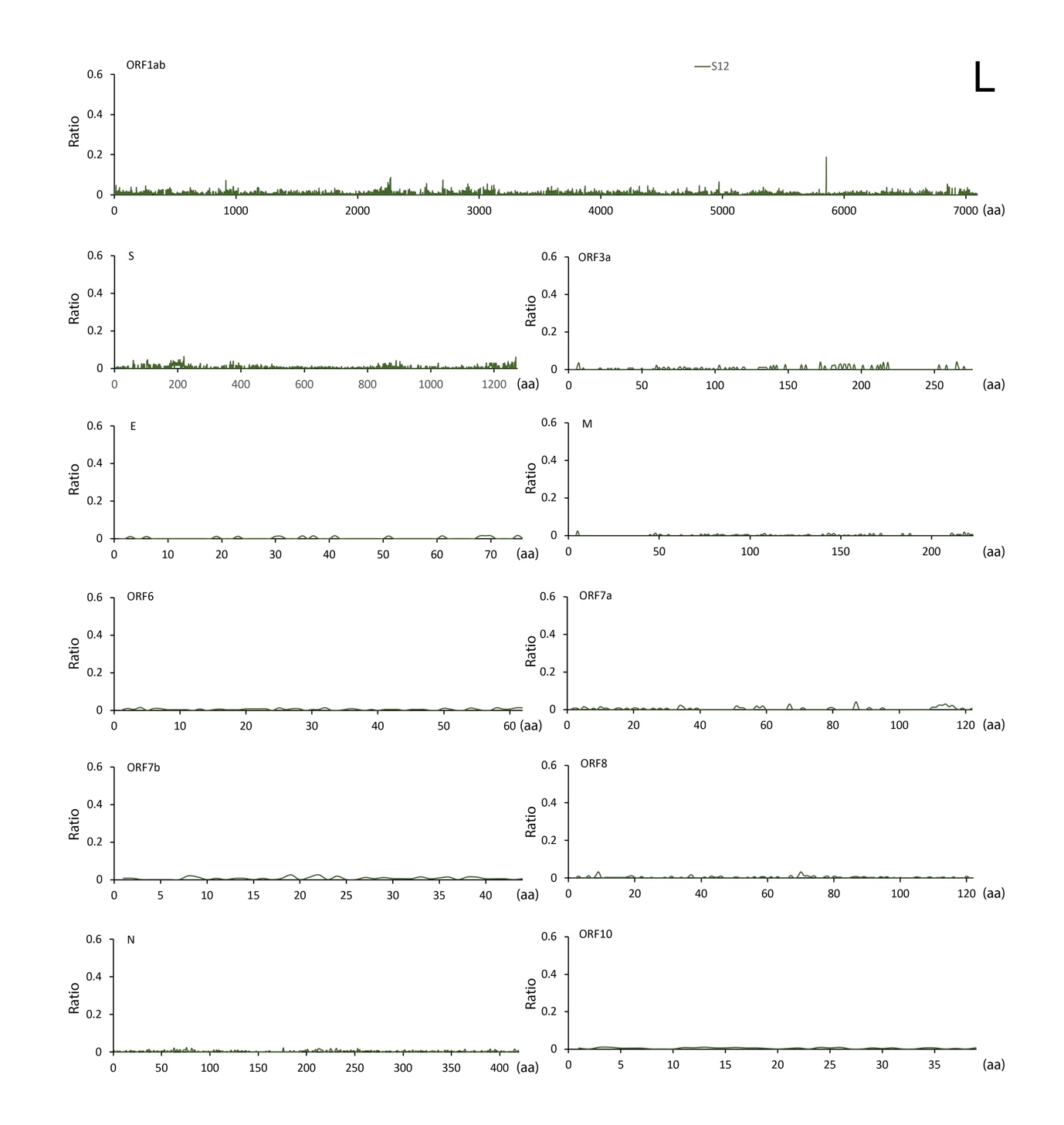

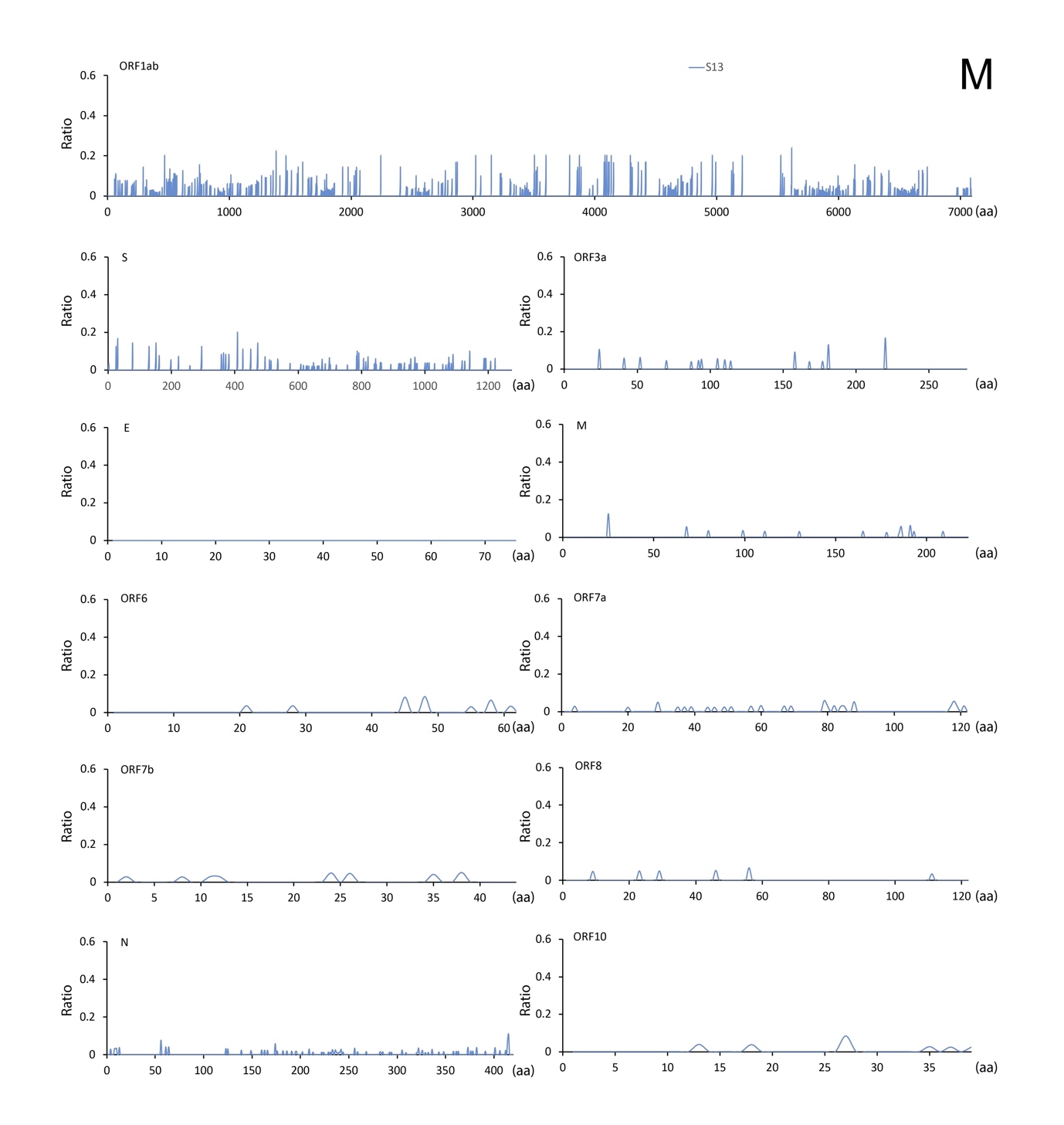

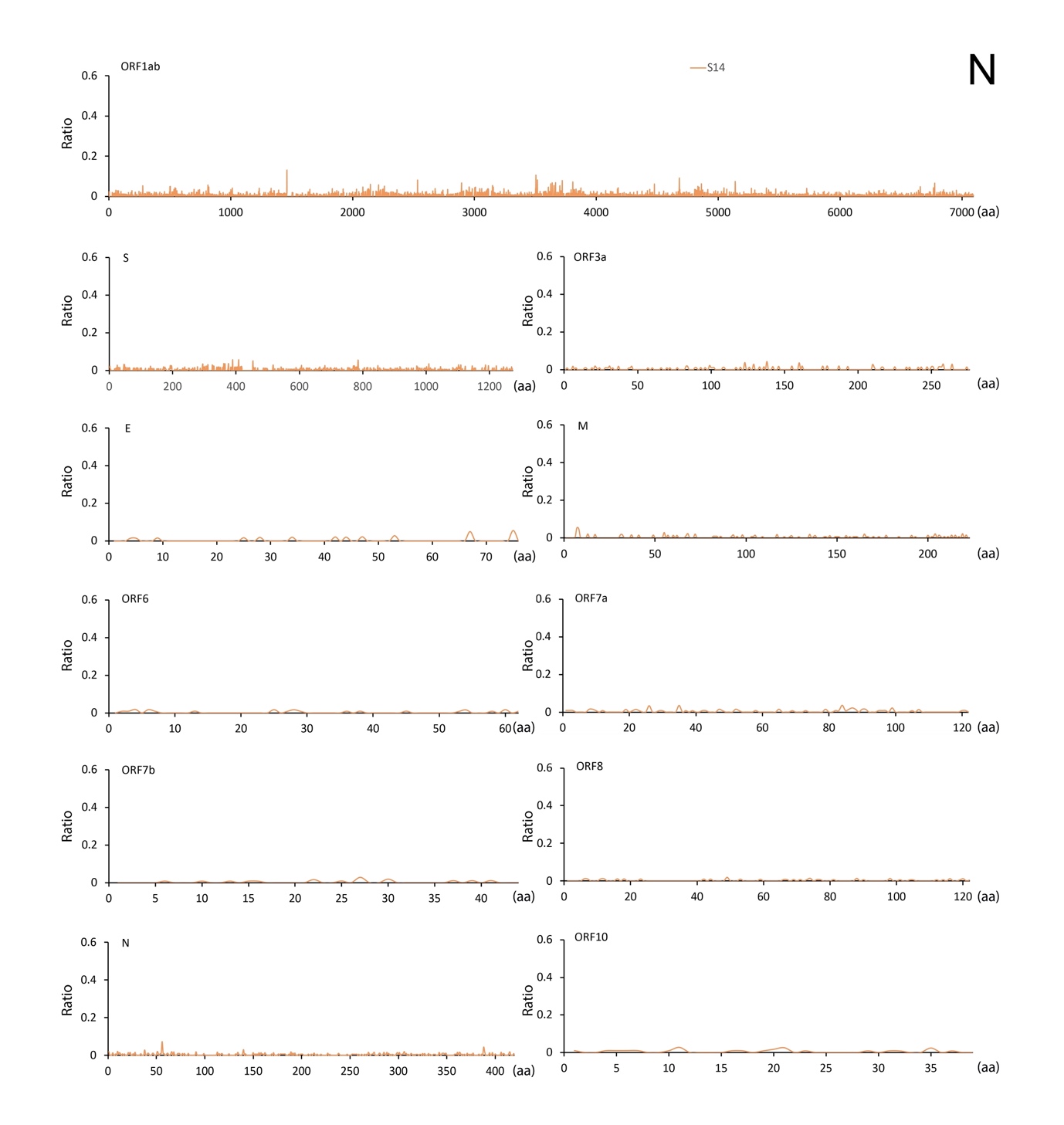

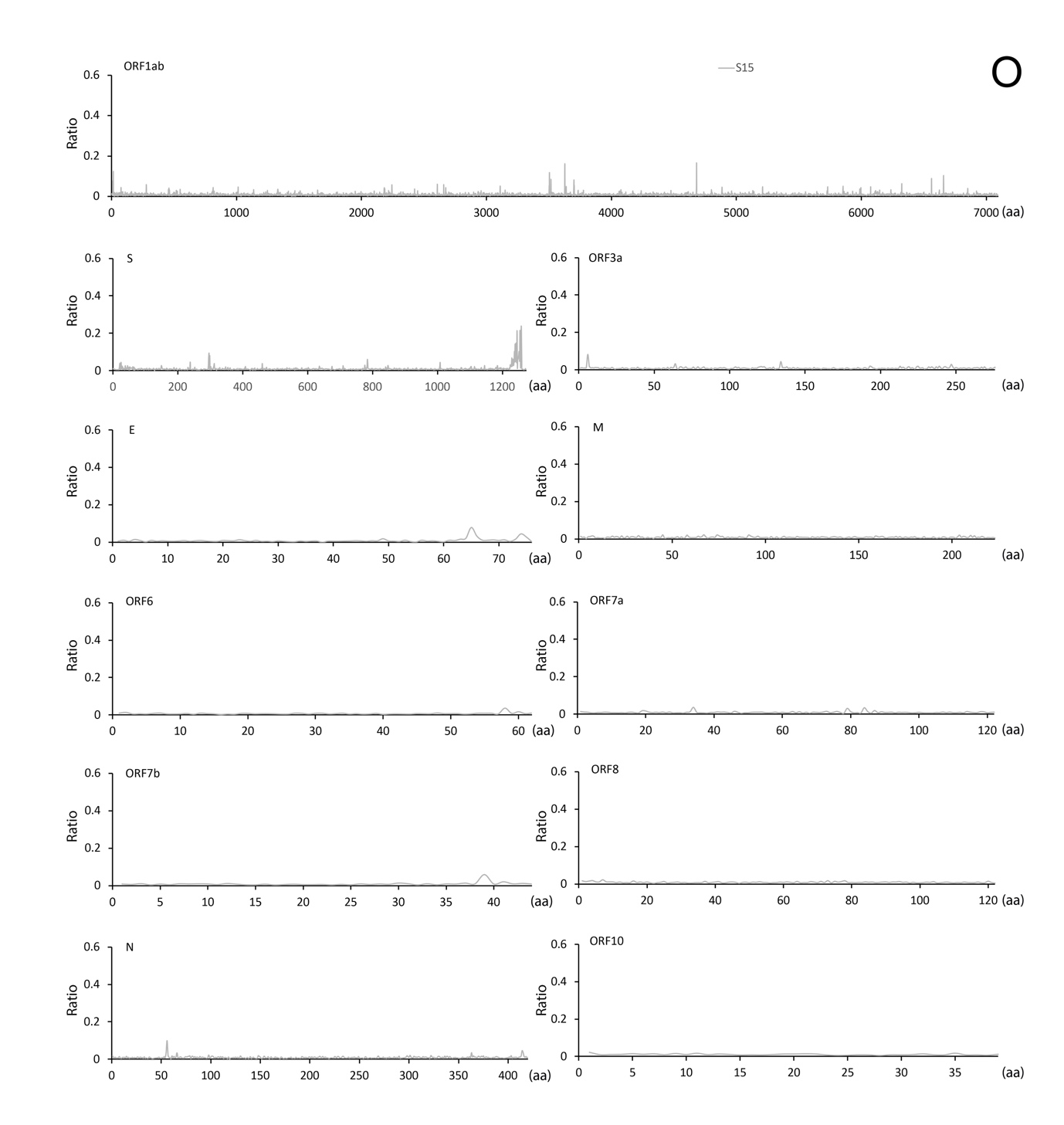

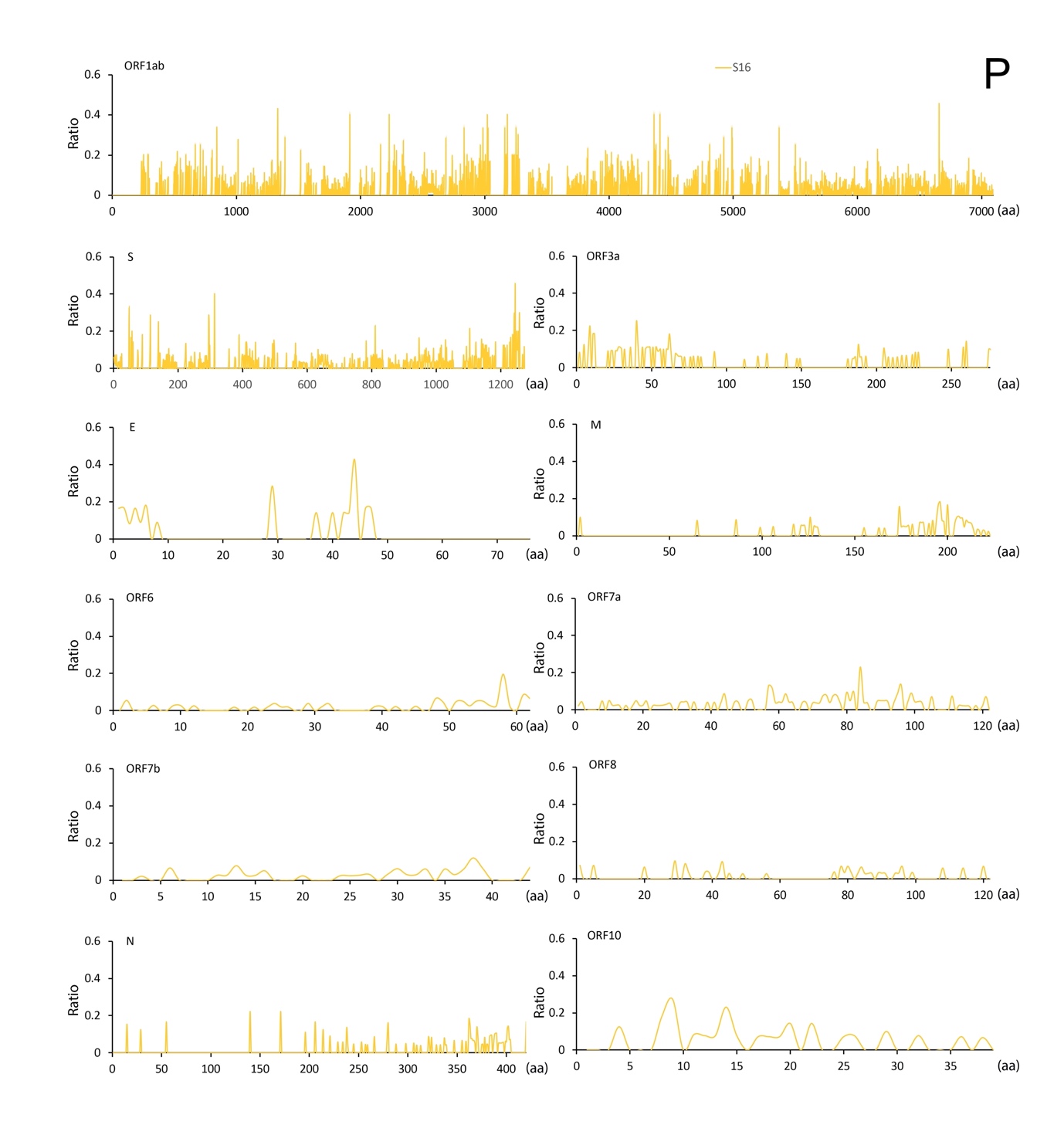
Fig. S1. Individual map of minor variant genomes across the SARS-CoV-2 genome for S1 (A), S2 (B), S3 (C), S4 (D), S5 (E), S6 (F), S7 (G), S8 (H), S9 (I), S10 (J), S11 (K), S12 (L), S13 (M), S14 (N), S15 (O), and S16 (P) in Fig. 1.

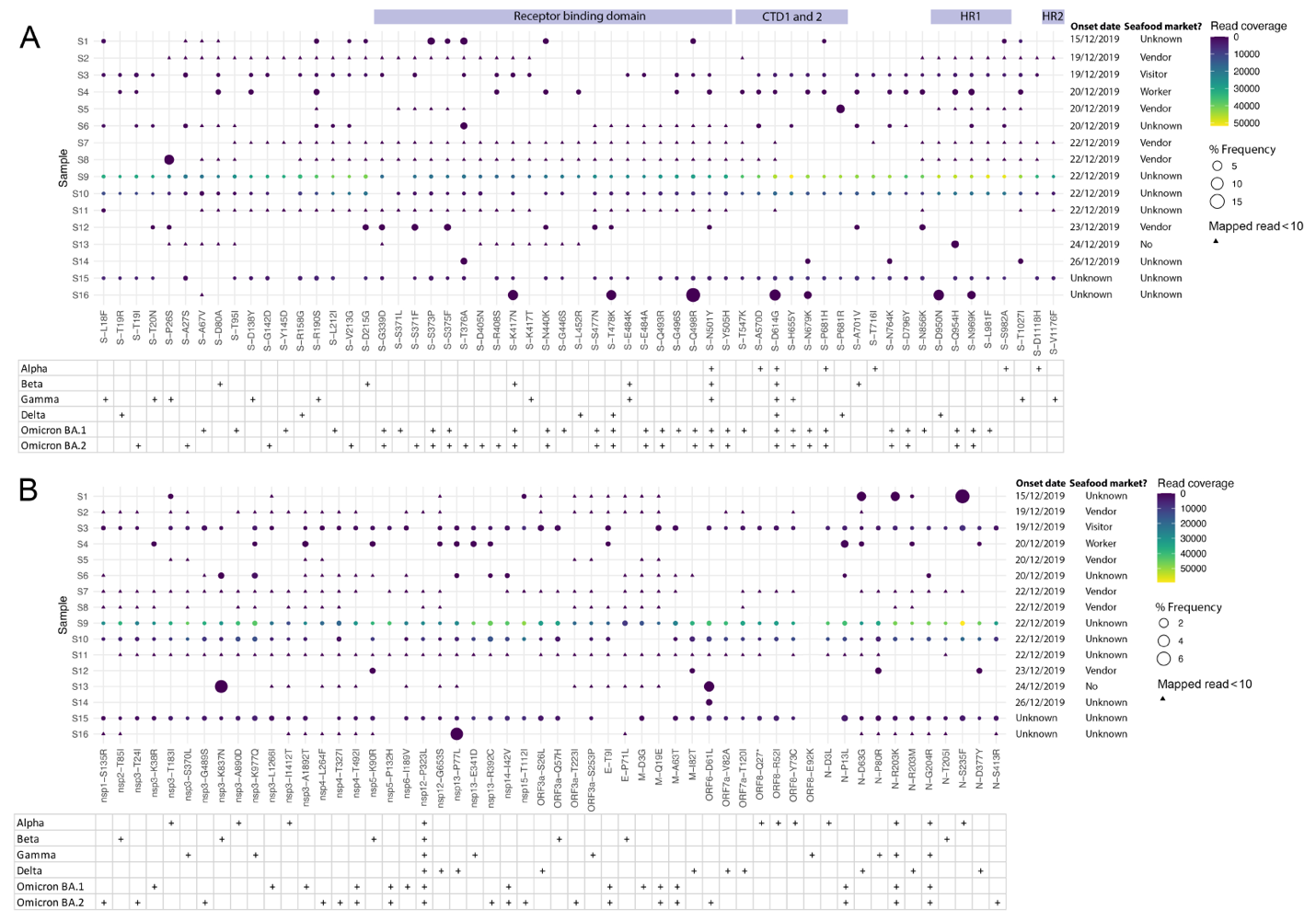

Fig. S2. Non-synonymous substitutions in the minor genomic variants of SARS-CoV-2 from each of the 16 down selected patients focusing on sites that define VoCs (https://covariants.org/variants) in the spike protein (A) and other regions of the genome (B). The amino acid site with coverage >= 10 were showed.

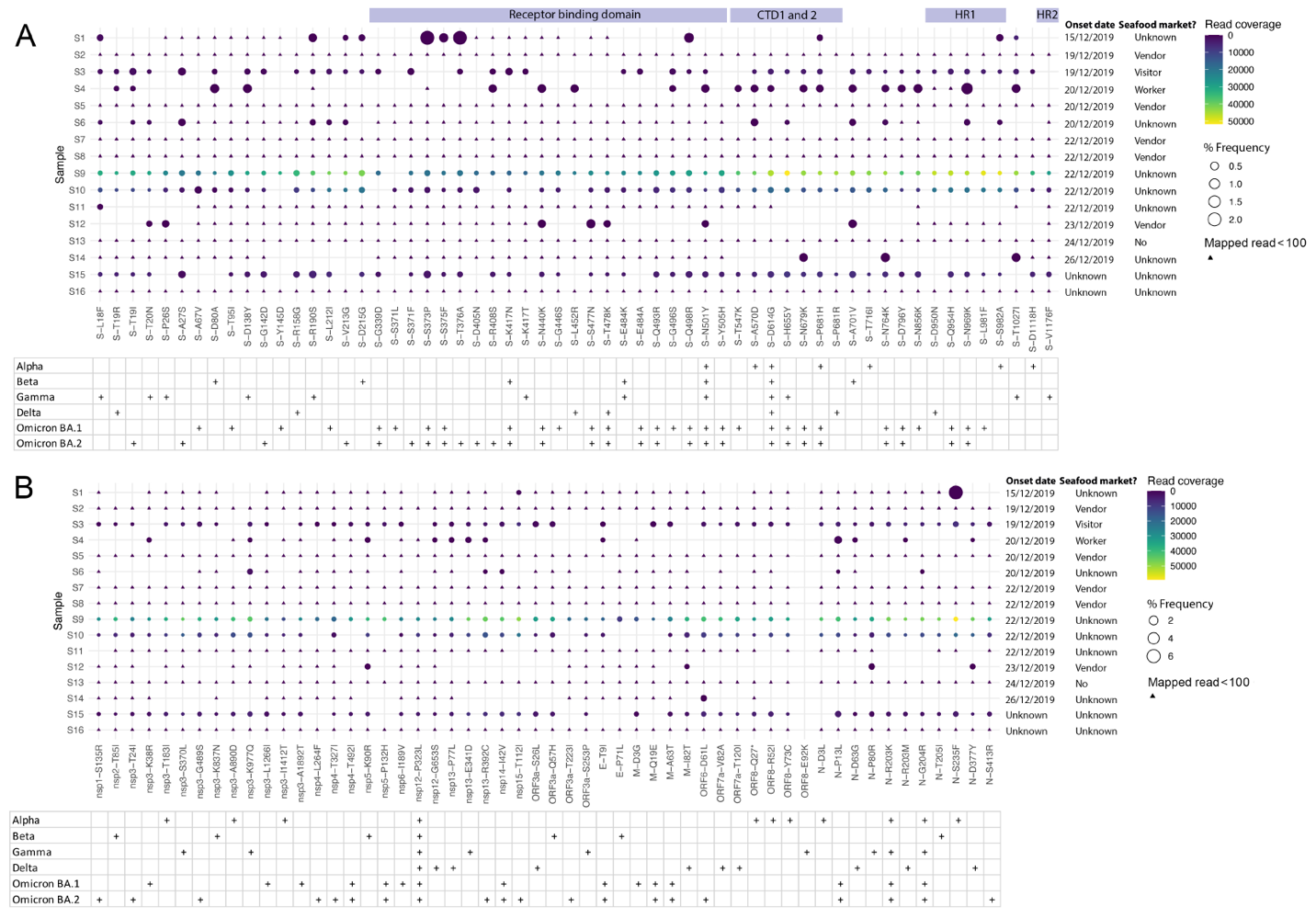

Fig. S3. Non-synonymous substitutions in the minor genomic variants of SARS-CoV-2 from each of the 16 down selected patients focusing on sites that define VoCs (https://covariants.org/variants) in the spike protein (A) and other regions of the genome (B). The amino acid site with coverage >= 100 were showed.

Table S1. Information of 16 sequenced samples collected from NCBI.

| **Sample ID** | **Accession ID** | **SRA ID** | **WHO ID** | **ID in article** | **Virus strain** | **lineage** | **Gender** | **Age** | **Onset date** | **Collection date** | **Wuhan seafood market** | **ICU** | **Sample** | **Sequencing method** | **Publication link** |
| --- | --- | --- | --- | --- | --- | --- | --- | --- | --- | --- | --- | --- | --- | --- | --- |
| S1 | SRX7705833 | SRR11059945 | - | - | nCov-RNA-3 | B | male | 40 | 15/12/2019 | 30/12/2019 | - | yes | BAL | Illumina HiSeq 2500 paired end sequencing | [https://doi.org/10.1093/cid/ciaa207](https://doi.org/10.1093/cid/ciaa203) |
| S2 | SRX7730880 | SRR11092063 | WHO_S04 | ICU-04 | WIV02 | B | male | 32 | 19/12/2019 | 30/12/2019 | Vendor | yes | BAL | Illumina HiSeq 3000 paired end sequencing | <https://www.nature.com/articles/s41586-020-2012-7> |
| S3 | SRX7705834 | SRR11059944 | - | - | nCov-RNA-4 | B | male | 61 | 19/12/2019 | 01/01/2020 | Visitor | yes | BAL | Illumina HiSeq 2500 paired end sequencing | <https://doi.org/10.1093/cid/ciaa203> |
| S4 | SRX7636886 | SRR10971381 | WHO_S06 | - | Hu-1 | B | male | 41 | 20/12/2019 | 26/12/2019 | Worker | - | BAL | Illumina MiniSeq paired end sequencing | <https://www.nature.com/articles/s41586-020-2008-3> |
| S5 | SRX7730884 | SRR11092059 | WHO_S08 | ICU-10 | WIV07 | B | male | 56 | 20/12/2019 | 30/12/2019 | Vendor | yes | BAL | Illumina HiSeq 3000 paired end sequencing | <https://www.nature.com/articles/s41586-020-2012-7> |
| S6 | SRX7705836 | SRR11059942 | - | - | nCov-RNA-6 | B | male | 56 | 20/12/2019 | 30/12/2019 | - | yes | BAL | Illumina HiSeq 2500 paired end sequencing | [https://doi.org/10.1093/cid/ciaa209](https://doi.org/10.1093/cid/ciaa203) |
| S7 | SRX7730882 | SRR11092061 | WHO_S11 | ICU-08 | WIV05 | B | female | 52 | 22/12/2019 | 30/12/2019 | Vendor | yes | BAL | Illumina HiSeq 3000 paired end sequencing | <https://www.nature.com/articles/s41586-020-2012-7> |
| S8 | SRX7730883 | SRR11092060 | WHO_S12 | ICU-09 | WIV06 | B | male | 40 | 22/12/2019 | 30/12/2019 | Vendor | yes | BAL | Illumina HiSeq 3000 paired end sequencing | <https://www.nature.com/articles/s41586-020-2012-7> |
| S9 | SRX7705831 | SRR11059947 | - | - | nCov-RNA-1 | B | female | 49 | 22/12/2019 | 30/12/2019 | - | no | BAL | Illumina HiSeq 2500 paired end sequencing | [https://doi.org/10.1093/cid/ciaa205](https://doi.org/10.1093/cid/ciaa203) |
| S10 | SRX7705832 | SRR11059946 | - | - | nCov-RNA-2 | B | female | 52 | 22/12/2019 | 30/12/2019 | - | yes | BAL | Illumina HiSeq 2500 paired end sequencing | [https://doi.org/10.1093/cid/ciaa206](https://doi.org/10.1093/cid/ciaa203) |
| S11 | SRX7705835 | SRR11059943 | - | - | nCov-RNA-5 | B | male | 40 | 22/12/2019 | 30/12/2019 | - | no | BAL | Illumina HiSeq 2500 paired end sequencing | [https://doi.org/10.1093/cid/ciaa208](https://doi.org/10.1093/cid/ciaa203) |
| S12 | SRX7730881 | SRR11092062 | WHO_S10 | ICU-06 | WIV04 | B | female | 49 | 23/12/2019 | 30/12/2019 | Vendor | yes | BAL | Illumina HiSeq 1000 paired end sequencing | <https://www.nature.com/articles/s41586-020-2012-7> |
| S13 | SRX7705837 | SRR11059941 | - | - | nCov-RNA-7 | B | female | 53 | 24/12/2019 | 01/01/2020 | No | no | BAL | Illumina HiSeq 2500 paired end sequencing | [https://doi.org/10.1093/cid/ciaa204](https://doi.org/10.1093/cid/ciaa203) |
| S14 | SRX7705838 | SRR11059940 | - | - | nCov-RNA-8 | B | male | 41 | 26/12/2019 | 30/12/2019 | - | no | BAL | Illumina HiSeq 2500 paired end sequencing | [https://doi.org/10.1093/cid/ciaa210](https://doi.org/10.1093/cid/ciaa203) |
| S15 | SRX8032203 | SRR11454614 | - | - | HBCDC-HB-01/2019 | B | female | 49 | - | 30/12/2019 | - | - | BAL | Illumina MiSeq paired end sequencing |  |
| S16 | SRX8032205 | SRR11454612 | - | - | HBCDC-HB-04/2019 | B | male | - | - | 30/12/2019 | - | - | sputum | Illumina MiSeq paired end sequencing |  |

BAL: bronchoalveolar lavage fluid.

Table S2. Nucleotide insertion and deletion called by freebayes.

| **Sample** | **Position** | **Reference** | **Alternative** | **Inserted nucleotide** | **Deleted nucleotide** | **Quality score** |
| --- | --- | --- | --- | --- | --- | --- |
| S1 | 8084 | GAAAAACT | GAAAACT | - | A | 4290.82 |
| S1 | 18976 | CAACACA | CAAACACA | A | - | 452.059 |
| S6 | 8837 | ATA | AA | - | T | 22.9267 |
| S6 | 13884 | ATA | AA | - | T | 43.2573 |
| S6 | 13893 | TTG | TATG | - | A | 21.4099 |
| S11 | 2550 | TAAACCAACCAT | TACCAACCAT | - | AA | 1276.76 |
| S11 | 6023 | TATCCAA | TATCAA | - | C | 1001.59 |
| S11 | 10024 | ACA | ATCA | T | - | 330.538 |
